# Centriole triplet microtubules are required for stable centriole formation and inheritance in human cells

**DOI:** 10.1101/147975

**Authors:** Jennifer T. Wang, Dong Kong, Christian R. Hoerner, Jadranka Loncarek, Tim Stearns

**Affiliations:** Department of Biology, Stanford University, Stanford CA; Laboratory of Protein Dynamics and Signaling, Center for Cancer Research – Frederick, National Cancer Institute, National Institutes of Health, Frederick MD; Department of Medicine – Division of Oncology, Stanford School of Medicine, Stanford CA; Department of Genetics, Stanford School of Medicine, Stanford CA

## Abstract

Centrioles are composed of long-lived microtubules arranged in nine triplets. In unicellular eukaryotes, loss of the noncanonical tubulins, delta-tubulin and epsilon tubulin, result in loss of the triplet microtubule structure. However, the contribution of triplet microtubules to mammalian centriole formation and stability is unknown. Here, we report the first characterization of delta-tubulin and epsilon-tubulin *null* human cells. Centrioles in cells lacking either delta-tubulin or epsilon-tubulin lack triplet microtubules and fail to undergo centriole maturation. These aberrant centrioles are formed *de novo* each cell cycle, but are unstable and do not persist to the next cell cycle, leading to a futile cycle of centriole formation and disintegration. Disintegration can be suppressed by paclitaxel treatment. Delta-tubulin and epsilon-tubulin physically interact, indicating that these tubulins act together to maintain triplet microtubules and that these are necessary for inheritance of centrioles from one cell cycle to the next.

## Introduction

The major microtubule organizing center of mammalian cells, the centrosome, is composed of a pair of centrioles with associated appendages and pericentriolar material. The centrioles have a nine-fold symmetry and are formed, in part, of long-lived microtubules, which persist through multiple cell divisions (Kochanski and Borisy, 1990; Balestra et al., 2015). In most organisms, including humans, the centriolar microtubules have a triplet structure, found only in centrioles. This structure consists of a complete A-tubule and associated partial B-tubule attached to the A-tubule wall, and a partial C-tubule attached to the B-tubule wall.

The molecular mechanisms involved in making triplet microtubules are not well-understood, even in the well-characterized somatic centriole cycle of mammalian cells. In these cells centrioles duplicate once per cycle, such that daughter cells receive exactly one pair of centrioles. Centriole duplication is initiated at the G1-S transition when the kinase PLK4 localizes to a single focus on the mother centriole (Sonnen et al., 2012). Subsequently, the cartwheel, formed by SASS6 oligomerization, assembles to template the 9-fold symmetry of the centriole (Guichard et al., 2017; Hilbert et al., 2016). Microtubules are added to the cartwheel underneath a cap of CP110 (Kleylein-Sohn et al., 2007). By G2-M, the triplet microtubules are completely formed (Vorobjev and Chentsov, 1982). Subsequently, the A- and B-tubules elongate to the full ~500 nm length of the centriole, forming a distal compartment with doublet microtubules and marked by POC5 (Azimzadeh et al., 2009). In mitosis, the cartwheel is lost, the newly-formed centriole becomes disengaged from its mother, and acquires pericentriolar material (Vorobjev and Chentsov, 1980; Vorobjev and Chentsov, 1982; Khodjakov and Rieder, 1999; Tsou and Stearns, 2006; Tsou et al., 2009). In G2-M of the following cell cycle, the centriole acquires appendages, marking its maturation into a centriole that can nucleate a cilium (Graser et al., 2007; Guarguaglini et al., 2005).

Members of the tubulin superfamily are critical for centriole formation and function. All eukaryotes have alpha-, beta- and gamma-tubulin, but the tubulin superfamily also includes three less-studied members, delta-tubulin, epsilon-tubulin, and zeta-tubulin. Recent work has shown that these noncanonical tubulins are evolutionarily co-conserved, making up the ZED tubulin module (Turk et al., 2015). In the unicellular eukaryotes Chlamydomonas, Tetrahymena, Paramecium and Trypanosoma, mutations in delta-tubulin or epsilon-tubulin result in centrioles that lack triplet microtubules (Dupuis-Williams et al., 2002; Dutcher and Trabuco, 1998; Dutcher et al., 2002; Gadelha et al., 2006; Garreau de Loubresse et al., 2001; Goodenough and StClair, 1975; Ross et al., 2013). Humans and other placental mammals have delta-tubulin and epsilon-tubulin, but lack zeta-tubulin (Findeisen et al., 2014; Turk et al., 2015). Here, we show that human cells lacking delta-tubulin or epsilon-tubulin also lack triplets, that this results in unstable centrioles and initiation of a futile cycle of centriole formation and disintegration, and identify an interaction between delta-tubulin and epsilon-tubulin.

## Results and Discussion

To determine the roles of delta-tubulin and epsilon-tubulin in the mammalian centriole cycle, null mutations in *TUBD1* and *TUBE1* were made using CRISPR/Cas9 genome editing in hTERT RPE-1 human cells. Recent work has established that loss of centrioles in mammalian cells results in a p53-dependent cell-cycle arrest (Bazzi and Anderson, 2014; Lambrus et al., 2015; Wong et al., 2015). We found that homozygous null mutations of delta-tubulin or epsilon-tubulin could only be isolated in *TP53* -/- cells, thus all subsequent experiments use RPE1 *TP53* -/- cells as the control.

Three *TUBD1 -/-* and two *TUBE1 -/-* cell lines were generated (Figure 1A and Figure 3 – supplemental figure 1). The *TUBD1 -/-* lines are all compound heterozygotes bearing, proximal to the cut site, small deletions of less than 20 base pairs on one chromosome and insertion of one base pair on the other, resulting in frameshift and premature stop mutations. The two *TUBE1 -/-* lines are compound heterozygotes bearing large deletions surrounding the cut site, that in each case remove an entire exon and surrounding DNA, including the ATG start site. In all cases, the next available ATG is not in-frame. We conclude that these alleles are likely to be null, or strong loss-of function mutations.

**Figure 1:**
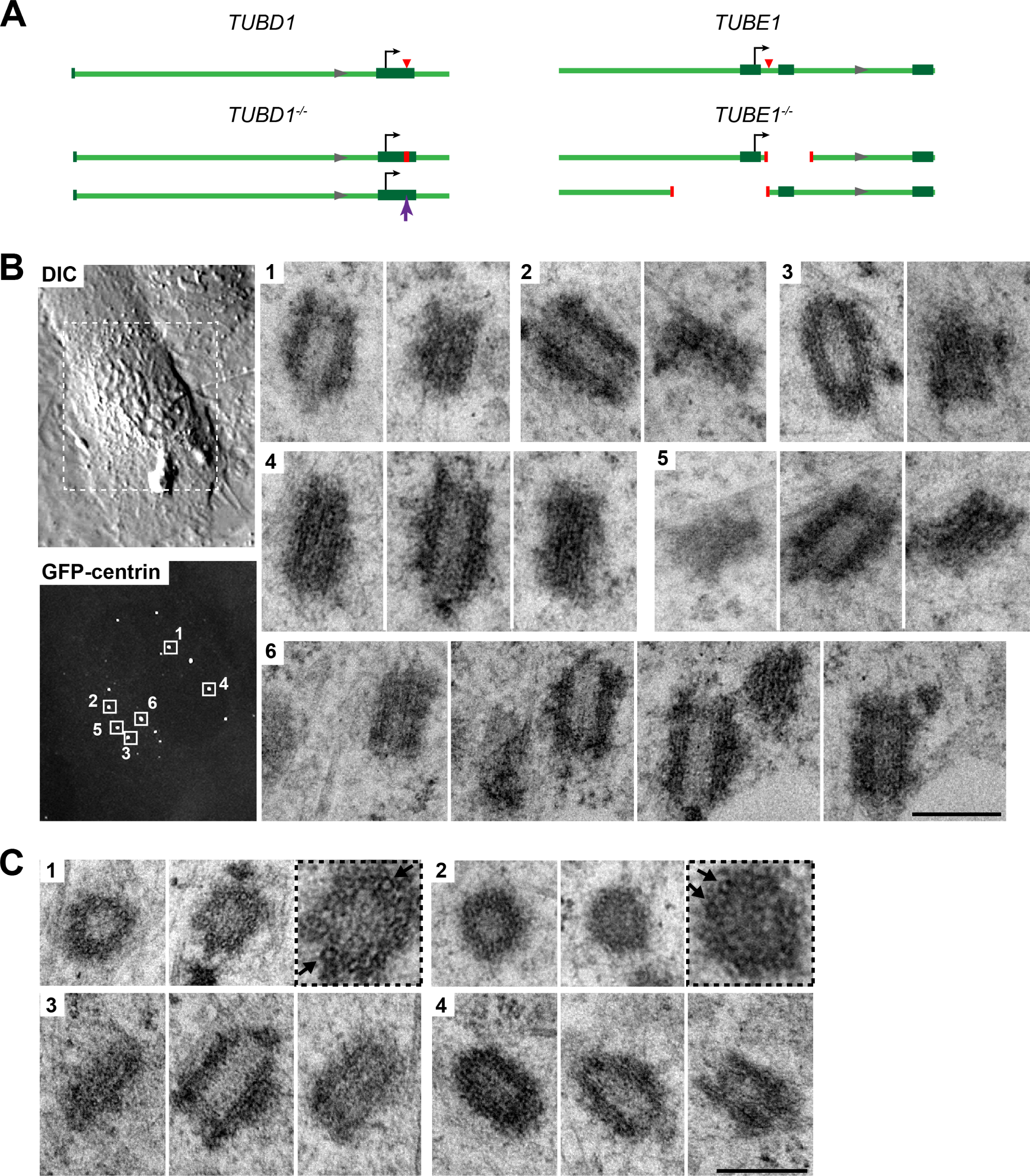
Centrioles in *TUBD1 -/-* and *TUBE1 -/-* cells are short and lack triplet microtubules. **A)** Gene loci for *TUBD1* (ch17:59889203-59891260) and *TUBE1* (ch6: 11207685- 11209742) in control and *TUBD1 -/-* and *TUBE1 -/-* cells (GRCh38.p7 Primary Assembly). Dark green boxes: exons, Arrows: translation start site, Red triangle: Cas9 cut site. *TUBD1 -/-* cells are compound heterozygotes, containing a 8 nt deletion (ch17: 59891019-59891026) on one allele, resulting in a frameshift and premature stop after 49 amino acids, and an insertion at nt 59891024 on the other, resulting in a frameshift and premature stop after 39 amino acids. The next ATG is not in-frame for either allele. *TUBE1 -/-* cells are also compound heterozygotes, containing a 266 nt deletion (ch6:112087525-112087790) on one allele, removing exon 2 and resulting in a frameshift and premature stop after 8 amino acids, and a 545 nt deletion (ch6:112086987-112087531) on the other allele, removing the first exon and the ATG. The next ATG is not in-frame for either allele. **B)** Centrioles from *TUBE1 -/-* cells. Left: DIC image and maximum intensity projection of *TUBE1 -/-* GFP-centrin cells. Boxed GFP-centrin foci were then analyzed by correlative electron microscopy. Right: electron micrographs of centrioles from boxed foci. Centrioles are numbered and serial sections are adjacent to each other. Scale bar: 250 nm **C)** Centrioles from *TUBD1 -/-* cells. Four centrioles are shown, and serial sections are adjacent to each other. For cross-sections from centrioles 1 and 2, a higher magnification image is placed third to show the presence of singlet microtubules (boxed). Arrows indicate singlet microtubules. Scale bar: 250 nm

We next assessed the phenotype of *TUBD1 -/-* and *TUBE1 -/-* cells stably expressing GFP-centrin as a marker of centrioles. Many cells in an asynchronous population had multiple, unpaired centrin foci (Fig. 1B). These foci also labeled with the centriolar proteins CP110 and SASS6 (see Figs 2 and 3). To determine whether these foci are centrioles, and to assess their ultrastructure, we analyzed them using correlative light-electron microscopy. In serial sections of interphase *TUBE1 -/-* (Fig 1 B) and *TUBD1 -/-* (Fig 1C) cells, some of the centrin-positive foci corresponded to structures that resemble centrioles, but were narrower and shorter than typical centrioles and lack appendages. They contained a ~80 nm wide central lumen, with luminal content resembling cartwheel architecture that extends throughout the centriole’s length (Fig 1 – supplement 1).

Centrioles in *TUBD1 -/-* and *TUBE1 -/-* cells were of similar diameter: 165 nm +/-15 nm in *TUBD1 -/-* cells (n = 19 centrioles), 164 nm +/-13 nm in *TUBE1 -/-* cells (n = 11 centrioles), compared to 220 nm diameter of the proximal end of typical mammalian cell centrioles (Loncarek et al., 2008; Wang et al., 2015). The reduced diameter of these aberrant centrioles is consistent with the presence of only singlet microtubules. Indeed, only singlet microtubules were identified in the two cross-sections observed, from *TUBD1 -/-* cells (Figure 1C). These results demonstrate that cells lacking either delta-tubulin or epsilon-tubulin form defective centrioles that lack normal triplet microtubules. This is similar to the defects reported for delta-tubulin and epsilon-tubulin mutants in unicellular eukaryotes.

Centrioles in both tubulin mutants were also shorter than typical, mature centrioles: 230 nm +/-45 nm in in *TUBD1 -/-* cells (n = 14 centrioles), and 271 nm +/-43 nm in in *TUBE1 -/-* cells (n = 11 centrioles), compared to approximately 500 nm for typical human cell centrioles (Paintrand et al., 1992). Newly-formed mammalian centrioles, or procentrioles, reach their full length by elongation in G2-M, creating a distal compartment that is a feature of centrioles in some, but not all, organisms. We sought to determine whether the aberrant centrioles in *TUBD1 -/-* and *TUBE1 -/-* cells are capable of elongation and formation of the distal compartment. We analyzed the ultrastructure of centrioles in a *TUBE1 -/-* prometaphase cell using correlative light-electron microscopy (Fig. 2A). These aberrant centrioles (n = 3) exhibited a striking morphological phenotype, consisting of two electron-dense segments, one of ~50 nm and the other of ~200 nm, connected by singlet microtubules spanning a gap of ~250 nm. The total length (~500 nm) of these structures approximates that of typical mature mammalian centrioles.

**Figure 2:**
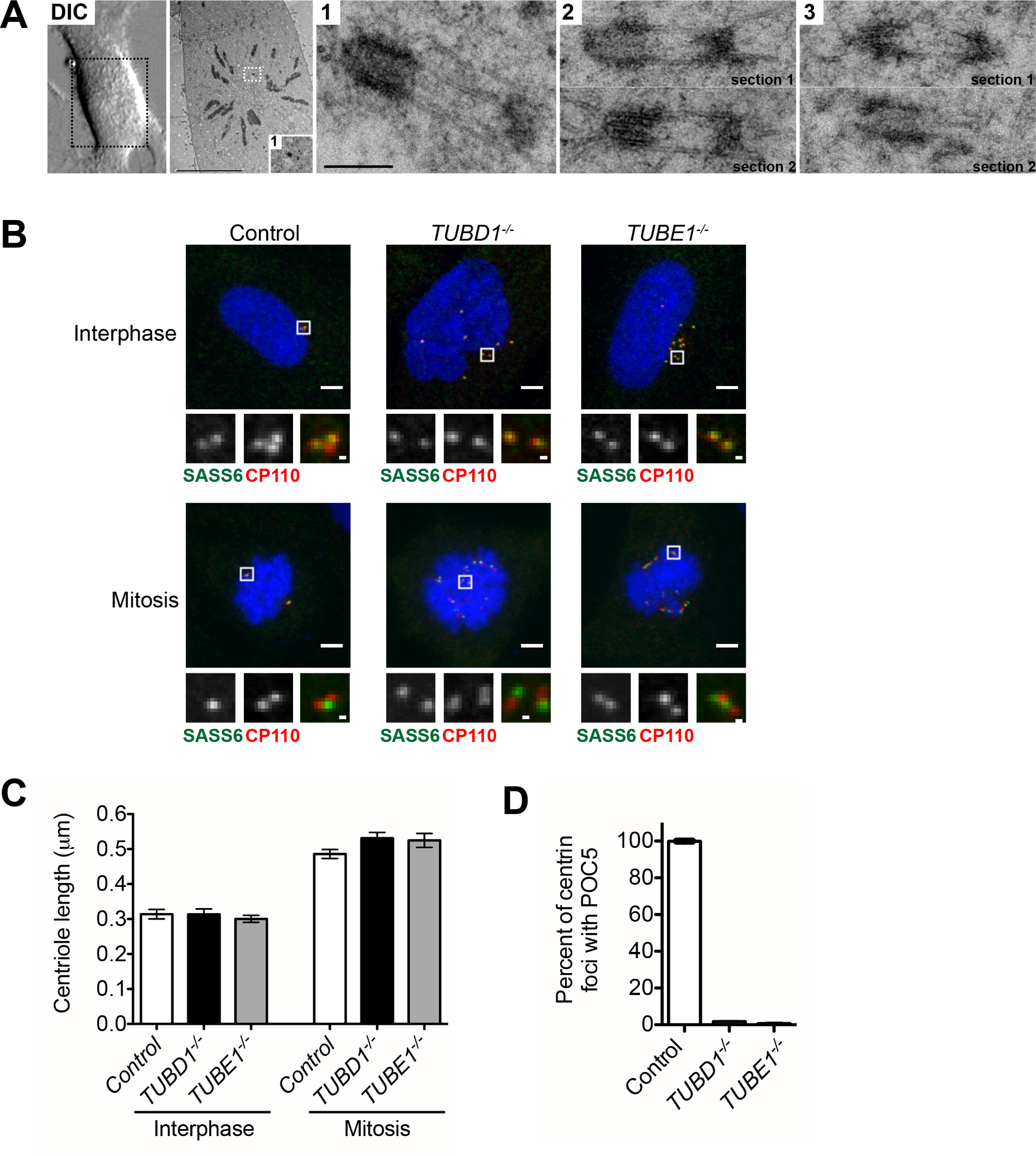
Centrioles in *TUBD1 -/-* and *TUBE1 -/-* cells elongate but fail to recruit POC5. **A)** Correlative light-electron micrographs of centrioles in a prometaphase *TUBE1 -/-* cell. Centrioles are numbered. Left: DIC image. Boxed centriole in overview corresponds to centriole 1. For centrioles 2 and 3, two serial sections are shown, as marked on figure. For each centriole, the proximal density is located on the left, and the distal density is located on the right. Scale bars: overview, 10 μm; inset: 250 nm. **B)** CP110 and SASS6 separation distance in interphase and mitotic cells. Images are maximum projections of 250 nm confocal stacks. Control cells are RPE-1 *TP53 -/-*. Scale bars: overview, 5 μm, inset: 500 nm. **C)** Quantification of CP110 and SASS6 separation distance. For each data point, 36 centrioles were measured. Control cells are RPE-1 *TP53 -/-*. Error bars represent the SEM. For each cell type, mitotic measurements are significantly different from interphase measurements (two-tailed unpaired t-test, p<0.0001). **D)** Quantification of the number of centrioles with POC5 localization in mitotic cells. Control cells are RPE-1 *TP53 -/-*. Bars represent the mean of three independent experiments with 200 centrioles each, error bars represent the SEM.

We hypothesized that the aberrant centrioles formed in *TUBD1 -/-* and *TUBE1 -/-* cells elongate in G2-M, but that only the A-tubule is present and thus able to elongate as a singlet. In this model, the shorter, distal density might correspond to the CP110 cap, under which the centriolar microtubules elongate (Kleylein-Sohn et al., 2007). The longer, 200 nm density corresponds to the proximal centriole end containing the cartwheel, as observed above in interphase cells. A prediction of this model is that the distance between CP110 and the SASS6 fluorescent labels would increase by about 200 nm in mitosis. We found that in *TUBD1 -/-* and *TUBE1 -/-* interphase cells, similar to control *TP53 -/-* cells, the centroids of CP110 and SASS6 foci were separated by a mean distance of 0.3 μm, whereas in mitotic cells the foci were separated by a mean distance of 0.5 μm (Fig 2B and 2C). Thus, centrioles in *TUBD1 -/-* and *TUBE1 -/-* cells elongate at the appropriate time in the cell cycle, and have a cap and proximal end typical of newly-formed centrioles. The lack of electron-dense structure between the cartwheel and cap might be due to a failure to recruit distal compartment components. Consistent with this, we found that the distal compartment component POC5 is absent from these aberrant centrioles (Fig 2D).

Together, these results indicate that the primary centriolar defect in cells lacking delta tubulin or epsilon-tubulin is the absence of triplet microtubules. To determine the consequences of loss of triplet microtubules on the centriole cycle and centrosome formation, we first determined the distribution of centrioles in asynchronously dividing cell populations, as determined by staining for the established centriole proteins centrin and CP110. *TP53 -/-* control cells had a typical centriole number distribution, with approximately 50% of cells having two centrioles, corresponding to cells in G1 phase, and 40% having three to four centrioles, corresponding to cells in S through M phases. In contrast, in *TUBD1 -/-* and *TUBE1 -/-* cells, approximately 50% of cells had 5 or more centriole foci, whereas 50% of cells had no detectable foci positive for both centrin and CP110 (Fig. 3A and 3B). Similar centriole distributions were found in other, independently derived, *TUBD1 -/-* and *TUBE1 -/-* cell lines. In addition, this phenotype could be rescued by expression of delta-tubulin and epsilon-tubulin, respectively (Fig 3 – supplement 1A – 1C).

**Figure 3:**
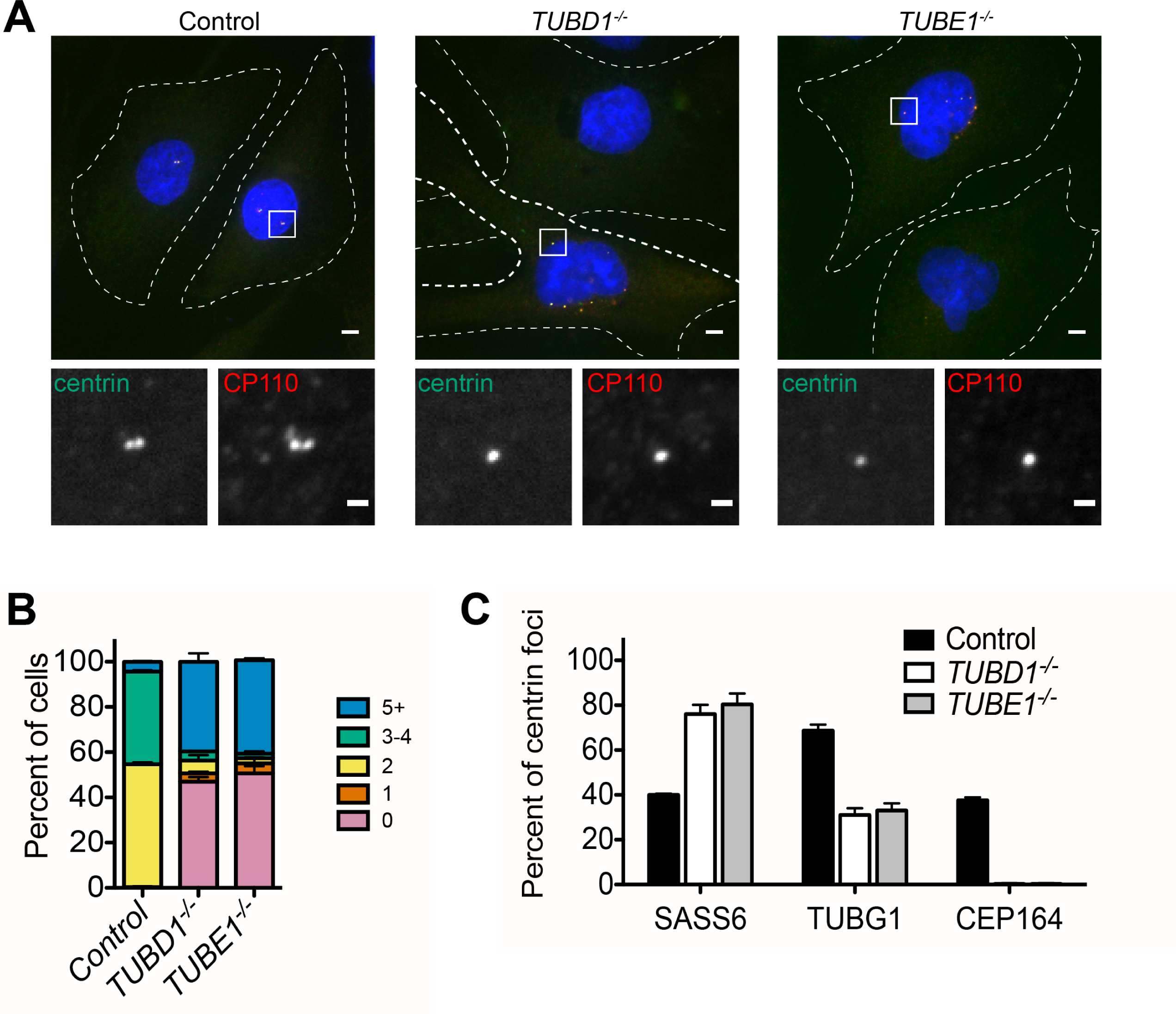
Centriole distribution and composition in *TUBD1 -/-* and *TUBE1 -/-* cells. **A)** Centriole phenotype for *TUBD1 -/-* and *TUBE1 -/-* cells. Two cells for each mutant are shown: one with no centrioles and the other with multiple centrioles. Control cells are RPE-1 *TP53 -/-*. Scale bars: overview, 5 μm; insets: 1 μm. **B)** Quantification of centriole number distribution in asynchronous cells, as measured by centrin and CP110 colocalization. Control cells are RPE-1 *TP53 -/-*. Bars represent the mean of three independent experiments with ≥100 cells each, error bars represent the SEM. **C)** Quantification of the percent of centrin foci that colocalize with indicated centriole markers. SASS6 marks early centrioles but is absent from mature centrioles, TUBG1 (gamma-tubulin) marks maturing centrioles, CEP164 marks completely mature centrioles. Control cells are RPE-1 *TP53 -/-*. Bars represent the mean of three independent experiments with ≥200 centrioles each, error bars represent the SEM.

We reasoned that a possible explanation for the centriole distribution in *TUBD1 -/-* and *TUBE1 -/-* cells is that the centriole structures we observed by EM are produced *de novo* in each cell cycle, and that these aberrant centrioles are unstable and do not persist into the next cell cycle. This hypothesis predicts that the aberrant centrioles in *TUBD1 -/-* and *TUBE1 -/-* cells would 1) not be paired, since de novo centrioles only form in the absence of an existing centriole, 2) lack markers of maturation such as distal appendages, since they would not persist to the point of acquiring such proteins, 3) fail to recruit substantial pericentriolar material, since the centriole-centrosome conversion occurs at entry to the next cell cycle, and 4) would be formed in S phase, and be lost at some point prior to the subsequent S phase.

In agreement with this hypothesis, the centrioles, as visualized by centrin and CP110 were never observed to be closely apposed, as is typical of wild-type cells (Fig. 3A). Rather, in interphase they appeared to be distributed within the central region of the cell (Fig. 3A). The centrioles in asynchronous *TUBD1 -/-* and *TUBE1 -/-* cells all lacked Cep164, a component of the centriolar distal appendage and marker of mature centrioles that have progressed through at least one cell cycle (Fig. 3C), whereas approximately 40% of all centrioles were positive for Cep164 in asynchronous control cells, consistent with the cycle of distal appendage acquisition (Nigg and Stearns, 2011). Lastly, most of the centrioles in *TUBD1 -/-* and *TUBE1 -/-* cells lacked detectable gamma-tubulin (Fig. 3C), and those that stained positive had less than centrioles in control cells (Fig 3 – supplement 1D). In addition, we noted that SASS6, the cartwheel protein that is present in nascent and recently-formed centrioles, but is lost from centrioles at the mitosis-interphase transition in human cells, was present in most of the centrioles in *TUBD1 -/-* and *TUBE1 -/-* cells, consistent with these centrioles originating in the observed cell cycle, but not having successfully persisted into the subsequent cell cycle.

To investigate the fate of newly-formed centrioles in *TUBD1 -/-* and *TUBE1 -/-* cells, we next tested the cell cycle-dependence of the formation and loss of aberrant centrioles in *TUBD1 -/-* and *TUBE1 -/-* cells (Fig 4A). As in previous experiments, about 50% of *TUBD1 -/-* and *TUBE1 -/-* cells in an asynchronous population had centrin and CP110- positive foci corresponding to aberrant centrioles. *TUBD1 -/-* and *TUBE1 -/-* cells were analyzed in different cell cycle stages as follows: G0/G1 – synchronized by serum withdrawal, S phase – identified from asynchronous culture by PCNA labeling, G2 – synchronized by the CDK1 inhibitor RO-3306, and M – identified from asynchronous culture by presence of condensed chromatin (Fig. 4A). *TUBD1 -/-* and *TUBE1 -/-* cells in G0/G1 mostly lacked centriole structures, whereas cells in S-phase, G2 and mitosis had them. These results indicate that in *TUBD1 -/-* and *TUBE1 -/-* cells, aberrant centrioles are formed in S-phase, persist into mitosis, and are absent in G1. We note that this loss of centriole structure is likely due to a specific event that occurs at the mitosis-interphase transition, rather than simply time since formation, since cells were arrested in G2 for 24 h, which is substantially longer than the normal progression through mitosis to G1, yet the centriole structures persisted (Fig 4A).

**Figure 4:**
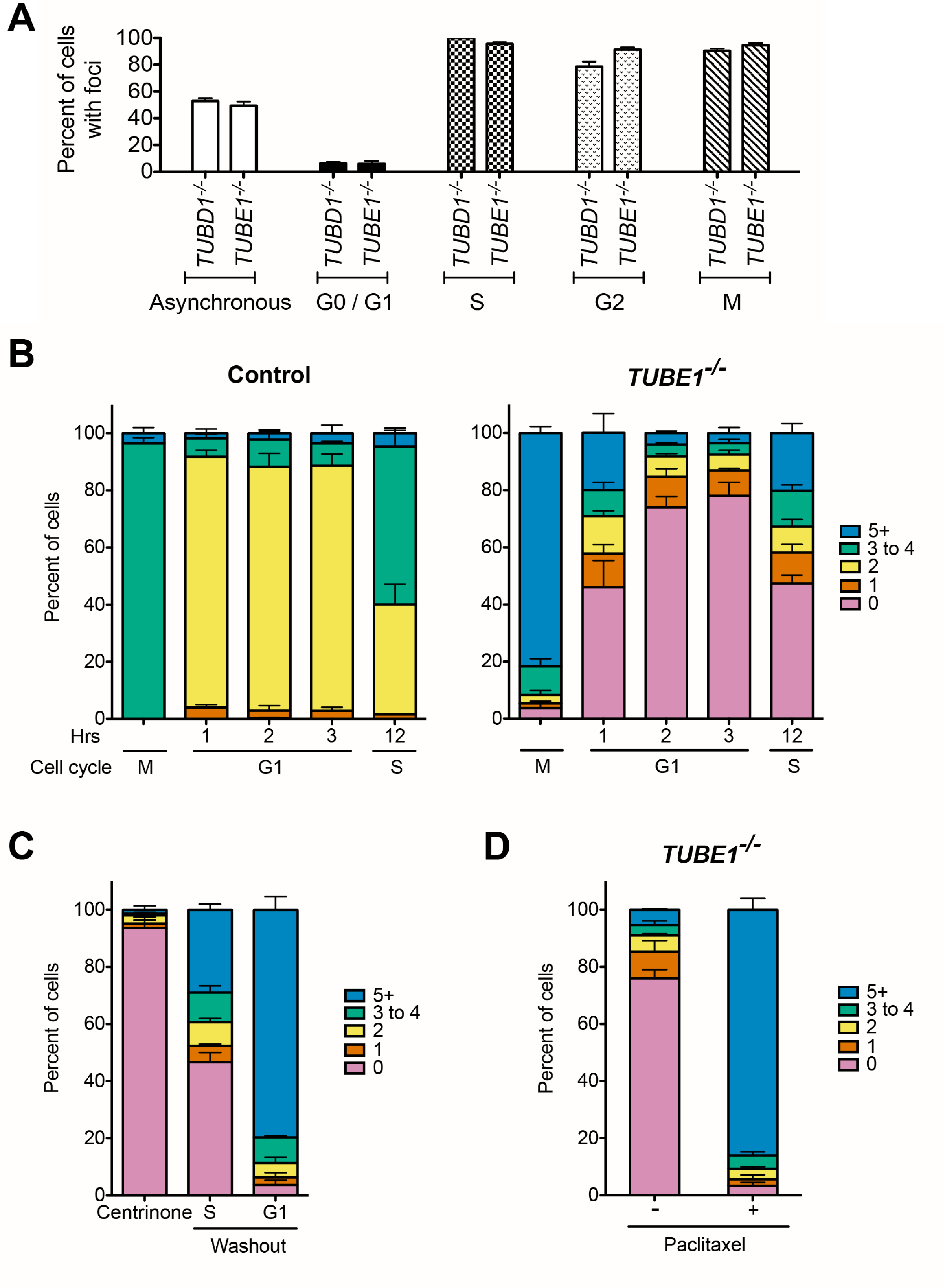
*TUBD1 -/-* and *TUBE1 -/-* cells undergo a futile centriole formation/disintegration cycle. **A)** Centriole presence in *TUBD1 -/-* and *TUBE1 -/-* cells is cell-cycle dependent. Quantification of the number of cells at each stage with centrin/CP1 1 0-positive centrioles. G0/G1 cells were obtained by serum withdrawal, S-phase by staining for PCNA, G2 by treatment with RO-3306, and mitosis by presence of condensed chromatin. Bars represent the mean of three independent experiments with ≥100 cells each, error bars represent the SEM. **B)** Quantification of the number of cells with centrin/CP110-positive centrioles at the indicated times after mitotic shakeoff. At 12 hours, 56 +/- 12% of TUBE1-/- cells entered S-phase, as marked by PCNA staining. Control cells are RPE-1 *TP53 -/-*. Bars represent the mean of three independent experiments with ≥150 cells each, error bars represent the SEM. **C)** *de novo* centrioles formed in the presence of TUBE1 do not disintegrate in G1. Quantification of the number of cells with centrin/CP110-positive centrioles after centrinone treatment. RPE1 *TP53 -/-* cells were treated with centrinone for at least 2 weeks, then centrinone was washed out from mitotic cells. Cells were analyzed in S-phase, 12 hours after washout when 36% of cells had entered S-phase, and in the following G1 after mitotic shakeoff. Bars represent the mean of three independent experiments with ≥100 cells each, error bars represent the SEM. **D)** Paclitaxel rescues centriole disintegration phenotype. *TUBE1 -/-* cells were either treated with paclitaxel or DMSO for 3 h in G2. Mitotic cells from both populations were harvested by mitotic shakeoff, and forced out of mitosis with RO-3306 for 3 h. Cells with micronuclei were analyzed for both conditions, and the percent of cells with indicated numbers of centrin/CP110-positive centrioles are shown. Bars represent the mean of three independent experiments with ≥100 cells each, error bars represent the SEM.

To more finely determine the timing of centriole loss in the mitosis-interphase transition, control or *TUBE1 -/-* cells were synchronized by mitotic shakeoff, and the presence of centriole foci was assessed over time as cells entered G1 (Fig 4B). In control cells, the number of centrioles follows the pattern expected from the centriole duplication cycle. In *TUBE1 -/-* cells, the majority of mitotic cells had centrioles. By 1 h after shakeoff, the fraction of interphase cells without centrioles had increased to 50%, and this fraction continued to increase at 2 h and 3 h after shakeoff. By 12 h after shakeoff, 56 +/-12% of cells had entered S-phase, and centriole structures began to appear, consistent with *de novo* centriole formation. Thus, delta-tubulin and epsilon-tubulin are not required to initiate centriole formation in human cells, but the aberrant centrioles that form in their absence are unstable and disintegrate during progression from M phase to the subsequent G1 phase.

Centrioles formed *de novo* can persist to form fully mature centrioles (Lambrus et al., 2015; Wong et al., 2015), but have also been reported to be structurally defective (Wang et al., 2015). We tested whether the phenotype we observed is specific to loss of delta-tubulin and epsilon-tubulin, rather than a property of *de novo* centrioles in general, by assessing whether *de novo* centrioles formed in the presence of delta-tubulin and epsilon-tubulin would also disintegrate upon cell cycle progression. RPE-1 *TP53 -/-* cells were treated with centrinone to inhibit PLK4 (Wong et al., 2015), a kinase required for centriole duplication, for more than 2 weeks to obtain acentriolar cells. Centrinone was then washed out from mitotic cells; by 12 h after shakeoff, 36% of cells had entered S-phase, and centriole structures began to appear, consistent with *de novo* centriole formation. However, in contrast to *TUBD1 -/-* and *TUBE1 -/-* cells, these *de novo* centrioles persisted through the subsequent G1 (Fig 4C). We conclude that centriole instability in *TUBD1 -/-* and *TUBE1 -/-* cells is due to a specific defect in their structure, and is not a general feature of *de novo* centrioles, similar to previous reports (La Terra et al., 2005).

We hypothesized that centriole disintegration may result from instability of the centriolar microtubules, perhaps as a result of elongation in G2-M phase. To test this, microtubules were stabilized in G2-M stage *TUBE1 -/-* cells by addition of the microtubule-stabilizing drug paclitaxel. This treatment did not inhibit centriole elongation (Fig 4 – supplement 1B). Cells were allowed to enter mitosis in the presence of paclitaxel, and subsequently forced out of mitosis using the CDK inhibitor RO-3306. This treatment was sufficient to stabilize centrioles from mutant cells in G1, compared with cells that had not been treated with paclitaxel (Fig 4D and Fig 4 – supplement 1). These stabilized centrioles lose their SASS6 cartwheel and fail to recruit detectable gamma-tubulin (Fig 4 – supplement 1). We conclude that stabilization of the centriolar microtubules in *TUBE1 -/-* cells stabilizes the centriole structure.

One striking observation of this work is that the phenotypes of delta-tubulin and epsilon tubulin null mutants are similar. This strongly suggests that the proteins work together to accomplish their function. To test this hypothesis, we assessed the ability of delta tubulin and epsilon-tubulin to interact by co-expression in human HEK293T cells. Epsilon-tubulin could be immunoprecipitated with delta-tubulin from co-expressing cells, and not from control cells (Fig 5A).

**Figure 5:**
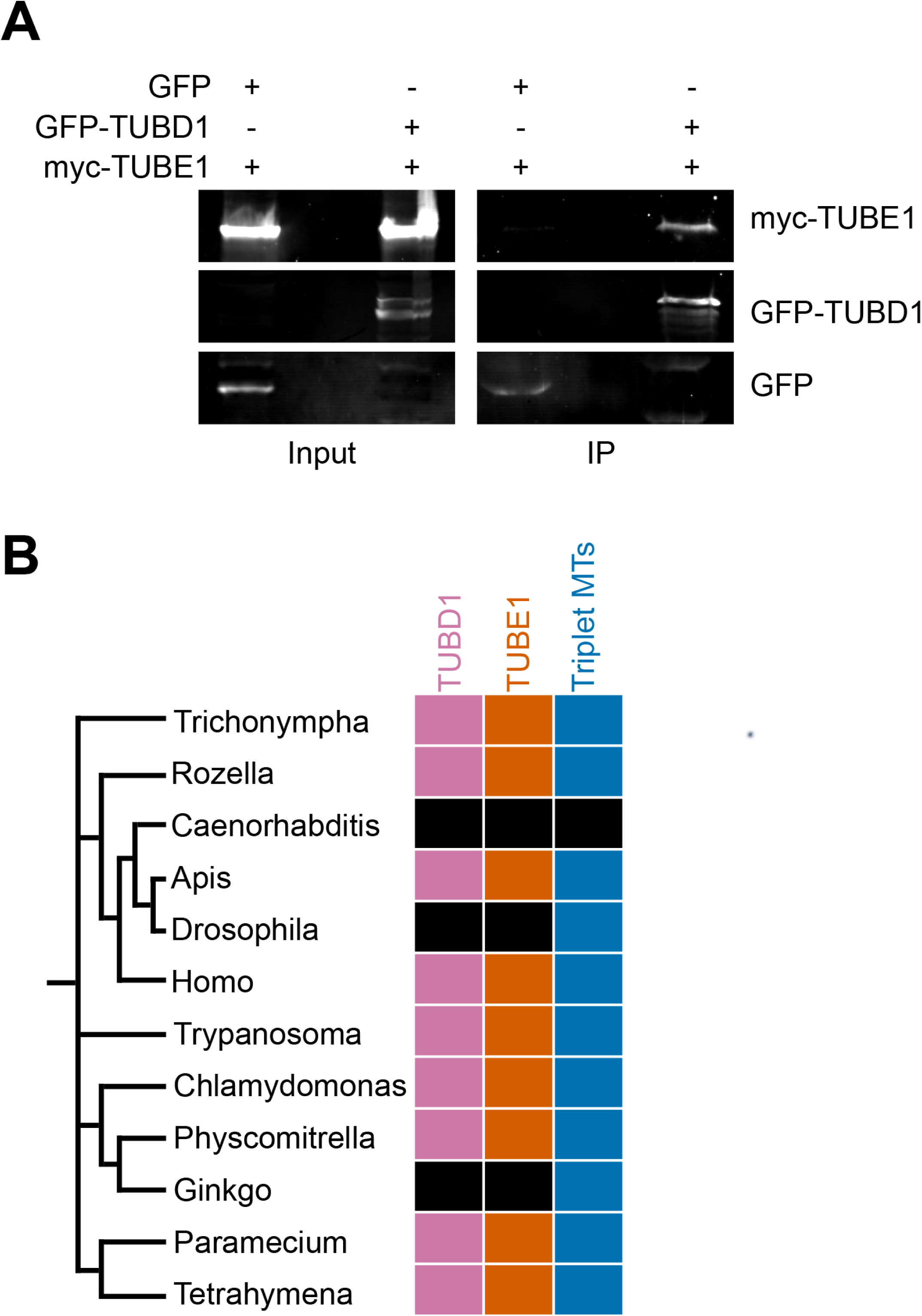
TUBD1 and TUBE1 interaction and evolutionary analysis. **A)** Co-immunoprecipitation of myc-TUBE1 and GFP-TUBD1. Complexes were immunoprecipitated (IP) with GFP-binding protein, and precipitated proteins were detected with anti-GFP and anti-Myc antibodies. **B)** Evolutionary analysis of the correlation of TUBD1 and TUBE1 presence with centriolar triplet microtubules, in organisms with centrioles. Black boxes represent genera in which the gene or feature is absent.

Together, our results show that delta-tubulin and epsilon-tubulin act together to create or stabilize structural features of centrioles. The most obvious such feature is the triplet microtubules, which define centrioles in most species, and are absent in delta-tubulin or epsilon-tubulin mutant cells in all organisms which have been examined. This suggests that delta-tubulin and epsilon-tubulin are required either to form the triplet microtubules, or to stabilize them against depolymerization. The former seems unlikely, since the presence of triplet centriolar microtubules is not strictly correlated with the presence of delta-tubulin and epsilon-tubulin in evolution (Fig 5B and Fig 5 - Supplemental Table 1). Among the organisms that completely lack the ZED tubulin module, *C. elegans* lacks triplet microtubules, but both Drosophila and the primitive plant *Ginkgo biloba* have triplet microtubules in their sperm cells. Since loss of the entire ZED tubulin module must have occurred independently in the dipteran insect and plant lineages, the most parsimonious interpretation is that triplet microtubule formation itself does not require delta-tubulin or epsilon-tubulin, rather than that these two lineages independently evolved mechanisms of triplet formation in their absence. Thus, we propose that delta-tubulin and epsilon-tubulin are required for stabilization of the centriolar triplets in most organisms, such that the centrioles can mature and recruit other proteins. We do not yet know the molecular basis of this differential requirement for delta-tubulin or epsilon-tubulins with respect to microtubule triplet stability. However, we note that the few centriole-bearing organisms that lack delta-tubulin and epsilon-tubulin have simpler centriole structures that lack distal appendages, and, to the extent it is possible to tell, lack a distal compartment that is typical of more complex centrioles.

Why do centrioles disintegrate in delta-tubulin and epsilon-tubulin mutant cells? We have shown that in *TUBD1 -/-* and *TUBE1 -/-* cells, aberrant centrioles with elongated singlet microtubules connecting the proximal and distal centriole segments become unstable as cells progress through mitosis. This is remarkably similar to the progressive loss of centrioles described in the original characterization of the epsilon-tubulin mutant *bald-2* by Goodenough and St. Clair (Goodenough and StClair, 1975). In human cells, Izquierdo, et al. reported that centrioles in *CEP295 -/-* human cells also become unstable upon cell cycle progression, due to a failure of centrioles to recruit pericentriolar material that coincides with loss of the cartwheel during the centriole-to centrosome conversion at the end of mitosis (Izquierdo et al., 2014). Although the phenotypes are outwardly similar to the phenotypes we describe here, CEP295 is conserved in species lacking delta-tubulin and epsilon-tubulin (Fu et al., 2015), and centrioles in Chlamydomonas do not undergo centriole-to-centrosome conversion. We propose that the post-duplication centriole elongation that creates the distal compartment of the centriole is a critical time in centriole stability, and that the triplet microtubules, either directly or through proteins that associate with them, are required to prevent centriole disassembly subsequent to that step. One possible basis for the instability is that events at the distal end of the centriole associated with preparing it to serve as a basal body for a cilium in G1 expose the ends of the centriolar microtubules. The doublet microtubules normally present at the end would be resistant to depolymerization in this model, but the singlets found in delta-tubulin and epsilon-tubulin mutants might be unstable. In accordance with this possibility, stabilization of centriolar microtubules with paclitaxel was able to prevent centriole disintegration, even when both the SASS6 cartwheel and pericentriolar material are lost (Fig 4D and Fig 4 – supplement 1). Another possibility is that centrioles lacking the normal triplet structure would likely also lack the A-C linker, which is visible in EM as a bridge between the A- and C-tubules of adjoining triplets. Perhaps the A-C linker is most important for stability after the full elongation of the centriolar microtubules. No components of the A-C linker have been identified, but the *poc1* mutant in Tetrahymena causes partial loss of this linker and results in instability of triplet microtubules (Meehl et al., 2016).

Here we have shown that delta-tubulin and epsilon-tubulin likely work together in a critical function for centriole structure and function, and that cells lacking delta-tubulin or epsilon-tubulin undergo a futile cycle of *de novo* centriole formation and disintegration. Our results show that in human cells, delta-tubulin and epsilon-tubulin act to stabilize centriole structures necessary for inheritance of centrioles from one cell cycle to the next, perhaps by stabilizing the main structural feature of centrioles, the triplet microtubules.

## Materials and Methods

### Cell culture

hTERT RPE-1 *TP53 -/-* cells were a gift from Meng-Fu Bryan Tsou (Memorial Sloan Kettering Cancer Center) and were cultured in DMEM/F-12 (Corning) supplemented with 10% Cosmic Calf Serum (CCS; HyClone). HEK293T cells were cultured in DMEM (Corning) supplemented with 10% CCS (HyClone). All cells were cultured at 37 °C under 5% CO2, and were routinely tested for mycoplasma contamination.

### Lentivirus production

Recombinant lentiviruses were made by cotransfection of HEK293T cells with the respective transfer vectors, second-generation lentiviral cassettes (packaging vector psPAX2 and envelope vector pMD2.G) using 1 μg/μL polyethylenimine (PEI; Polysciences). The medium was changed 6-8 h after transfection, and viral supernatant was harvested after an additional 48 h.

### Generation of *TUBD1 -/-* and *TUBE1 -/-* cells and rescue lines

hTERT RPE1 *TP53 -/-* GFP-centrin cells were made by transduction with mEGFP centrin2 lentivirus and 8 μg/mL Sequabrene carrier (Sigma-Aldrich). Cells were cloned by limiting dilution into 96-well plates.

*TUBD1 -/-* cell lines were generated using lentiCRISPRv2 (Addgene plasmid #52961 (Sanjana et al., 2014; Shalem et al., 2014) with the sgRNA sequence CTGCTCTATGAGAGAGAATG. hTERT RPE1 *TP53 -/-* GFP-centrin cells were transduced with lentivirus and 8 μg/mL Sequabrene for 72 hours, then passaged into medium containing 6 μg/mL puromycin. Puromycin-containing culture medium was replaced daily for 5 days until all cells in uninfected control had died. Puromycin-resistant cells were cloned by limiting dilution into 96-well plates, followed by genotyping and phenotypic analysis.

*TUBE1 -/-* cell line 1 was generated using pX330 (Addgene plasmid #42230 Cong et al., 2013) with the sgRNA sequence GGGTAGAGACCTGGTCGCCG (pX330-TUBE1). hTERT RPE-1 *TP53 -/-* cells were transiently co-transfected with pX330-TUBE1 and EGFP-expressing vector pEGFP-N1 (Clontech) at 9:1 ratio using Continuum Transfection Reagent (Gemini Bio-Products). GFP-positive cells were clonally sorted into single wells of 96-well plates by FACS, followed by genotyping and phenotypic analysis. Cells were subsequently transduced with GFP-centrin2 lentivirus for CLEM. *TUBE1 -/-* cell line 2 was generated using lentiCRISPRv2 with the sgRNA sequence GCGCACCACCATGACCCAGT. Transduction and selection were carried out as for *TUBD1 -/-* cell lines.

Both rescue construct transfer vectors contained opposite orientation promoters: EF-1alpha promoter driving monomeric Kusabira Orange kappa (mKOk) with rabbit beta-globin 3’UTR, as well as mouse PGK promoter driving the rescue construct with WPRE. For the delta-tubulin rescue construct, silent mutations were made in the PAM and surrounding sequence such that it was no longer complementary to the lentiCRISPR sgRNA (C117G and A120T) using QuikChange Lightning Site-Directed Mutagenesis Kit (Agilent). For the epsilon-tubulin rescue construct, full-length *TUBE1* cDNA was used. Using these transfer vectors, lentivirus was produced and *TUBD1 -/-* and *TUBE1 -/-* cells, respectively, were transduced. For rescue experiments, cells expressing mKOk were counted.

### Correlative light and electron microscopy

Correlative light and electron microscopy (CLEM) was performed as described previously (Kong and Loncarek, 2015), using hTERT RPE-1 *TP53 -/- TUBD1 -/-* and *TP53 -/- TUBE1 -/-* GFP-centrin cells. Cells in Rose chambers were enclosed in an environmental chamber at 37 °C and imaged on an inverted microscope (Eclipse Ti; Nikon, Tokyo, Japan) equipped with a spinning-disk confocal head (CSUX Spinning Disk; Yokogawa Electric Corporation, Tokyo, Japan). After analysis by live imaging, Rose chambers were perfused with freshly prepared 2.5% glutaraldehyde, and 200-nm thick *Z*-sections spanning the entire cell were recorded to register the position of centrioles. Cell positions on coverslips were then marked by diamond scribe. Rose chambers were disassembled, and cells were washed in PBS, followed by staining with 2% osmium tetroxide and 1% uranyl acetate. Samples were dehydrated and embedded in Embed 812 resin. The same cells identified by light microscopy were then serially sectioned. The 80 nm-thick serial sections were transferred onto copper slot grids, stained with uranyl acetate and lead citrate, and imaged using a transmission electron microscope (H-7650; Hitachi, Tokyo, Japan).

### Immunofluorescence

Cells were grown on poly-L-lysine-coated #1.5 glass coverslips. Cells were washed with PBS, then fixed with -20 °C methanol for 15 min. Coverslips were then washed with PBS and blocked with PBS-BT (3% BSA, 0.1% Triton X-100, 0.02% sodium azide in PBS) for 30 min. Coverslips were incubated with primary antibodies diluted in PBS-BT for 1 h, washed with PBS-BT, incubated with secondary antibodies and DAPI diluted in PBS-BT for 1 h, then washed again. Samples were mounted using Mowiol (Polysciences) in glycerol containing 1,4,-diazobicycli-[2.2.2]octane (DABCO, Sigma-Aldrich) antifade.

### Antibodies

Primary antibodies used for immunofluorescence: mouse IgG2b anti centrin3, clone 3e6 (1:1000, Novus Biological), mouse IgG2a anti centrin, clone 20H5 (1:200, EMD Millipore), rabbit anti CP110 (1:200, Proteintech), mouse IgG2b anti SASS6 (1:200, Santa Cruz), mouse IgG1 anti gamma-tubulin, clone GTU-88 (1:1000, Sigma-Aldrich), rabbit anti POC5 (1:500, Bethyl Laboratories), rabbit anti CEP164 (1:500, described previously (Lee et al., 2014), mouse IgG2a anti PCNA (1:500, BioLegend). Primary antibodies used for Western blotting: goat anti GFP (1:500, Rockland), mouse IgG1 anti myc, clone 9e10 (1:100, Developmental Studies Hybridoma Bank). For immunofluorescence, AlexaFluor conjugated secondary antibodies (Thermo-Fisher) were diluted 1:1000. For Western blotting, IRDye conjugated donkey secondary antibodies (LiCOR) were diluted 1:20,000.

### Drug treatments and mitotic shakeoff

For cell cycle analyses, *TUBD1 -/-* or *TUBE1 -/-* cells were seeded onto coverslips, then synchronized in G0/G1 by serum withdrawal for 24 h, or in G2 with 10 μM RO-3306 for 24 h. Cells were fixed for immunofluorescence and analyzed for centrin/CP110 presence. Mitotic shakeoff was performed on asynchronously growing cells. One pre-shake was performed to improve synchronization. Cells were fixed at indicated times and analyzed for centrin/CP110 presence. For centrinone experiments, hTERT RPE-1 *p53 -/-* cells were treated with 125 nM centrinone for ≥ 2 weeks, and centrinone-containing medium was replaced on top of cells daily. For centrinone washout, cells were washed twice with PBS, then mitotic shakeoff was performed with centrinone-free medium. A subset of cells were fixed for immunofluorescence 12 h after shakeoff, when cells had entered S-phase. 19 h after shakeoff, a second shakeoff was performed to harvest cells that entered mitosis. Cells were fixed 3 h post-second shakeoff for immunofluorescence, and analyzed for centrin/CP110 presence. For paclitaxel experiments, mitotic cells were removed by shakeoff from an asynchronous population, then 15 μM paclitaxel or DMSO was added to the cells remaining on the dish. For both populations, G2-phase cells were allowed to enter mitosis, and then harvested in mitosis by shakeoff 3 h later. Cells were plated on coverslips and forced to exit mitosis by treatment with 10 μM RO-3306, then fixed for immunofluorescence 3 h later. Cells with micronuclei were analyzed for centrin/CP110 presence in both conditions.

### Immunoprecipitation

HEK293T cells were co-transfected with GFP-delta-tubulin and myc-epsilon-tubulin, or GFP and myc-epsilon-tubulin using PEI. 48 hours after transfection, cells were harvested and lysed in lysis buffer (50 mM Hepes pH7.4, 150 mM NaCl, 1 mM DTT, 1 mM EGTA, 1 mM MgCl2, 0.25 mM GTP, 0.5% Triton X-100, 1 μg/ml each leupeptin, pepstatin, and chymostatin, and 1 mM phenylmethylsulfonyl fluoride). Insoluble material was pelleted, and soluble material was incubated at 4 °C with GFP-binding protein (Rothbauer et al., 2008) coupled to NHS-activated Sepharose 4 Fast Flow resin (GE Healthcare) for 2 h. Beads were pelleted at 500 g for 1 min, washed three times with lysis buffer, then eluted in sample buffer and the eluate was run on SDS-PAGE gels. Western blots were scanned on a LiCOR imager and analyzed using ImageJ.

#### Acknowledgements

We thank Meng-Fu Bryan Tsou for the gift of hTERT RPE-1 *TP53 -/-* cells, Olga Cormier for help with evolutionary analysis, and David Breslow and Max Nachury for sharing unpublished data. This work was supported by National Research Service Award grant 5 F32 GM117678 to J.T.W., the Intramural Research Program of the National Institutes of Health, National Cancer Institute, Center for Cancer Research to J.L., and NIH grant R01GM052022 to T.S.

---

**Figure 1 – figure supplement 1:**
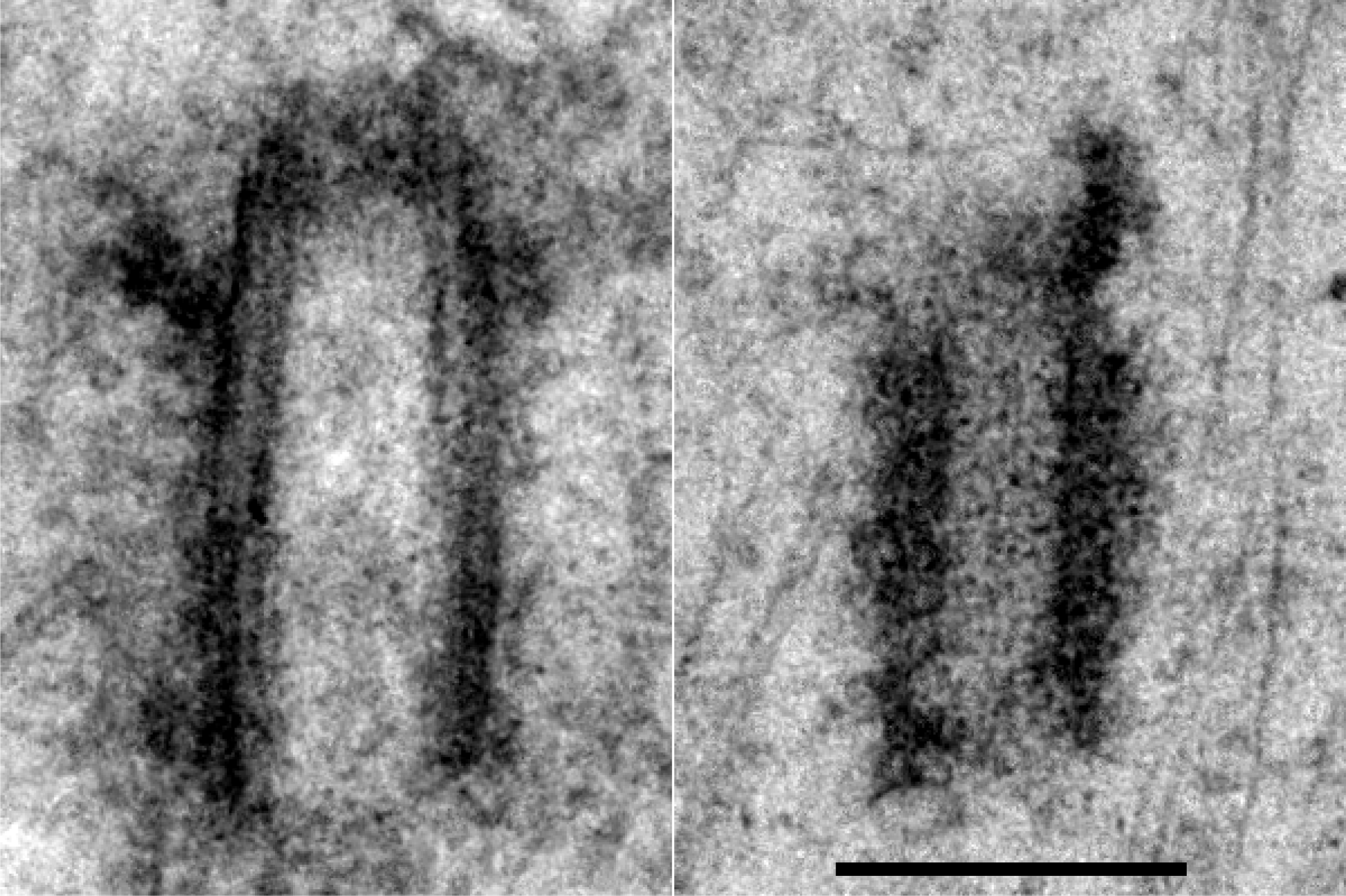
Comparison to control centrioles. Left: mature mother centriole from wildtype RPE-1 cells. Right: centriole from *TUBE1 -/-* cell. Scale bar: 250 nm

**Figure 3 – figure supplement 1:**
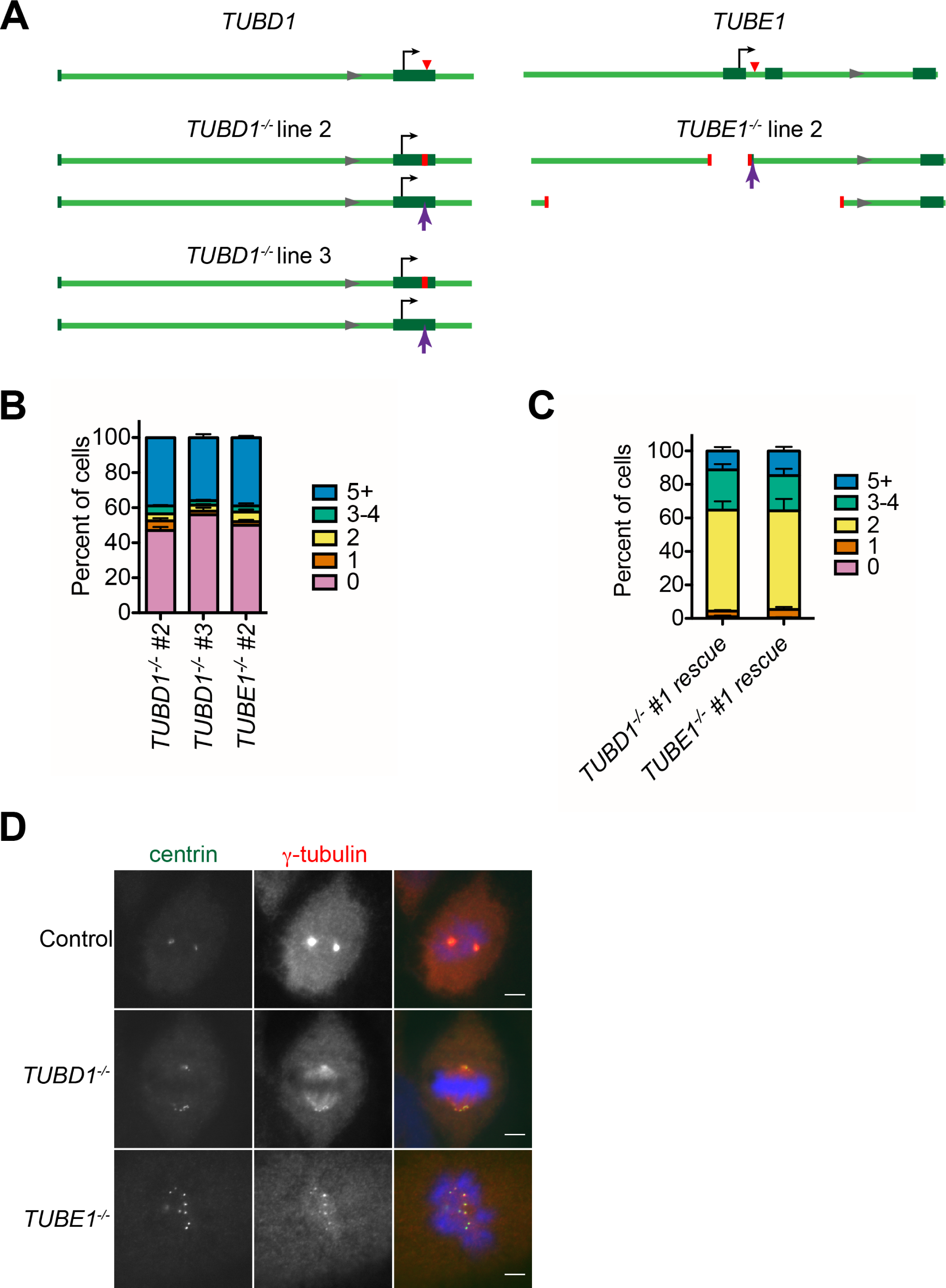
Centriole distribution in independently-derived clonal cell lines and rescue of the phenotype. **A)** Gene loci for TUBD1 (ch17:59889203-59891260) and TUBE1 (ch6: 11207685- 11209742) in control and *TUBD1 -/-* and *TUBE1 -/-* cells (GRCh38.p7 Primary Assembly). Dark green boxes: exons, Arrows: translation start site, Red triangle: Cas9 cut site. *TUBD1 -/-* line 2 is a compound heterozygote, containing a 4 nt deletion (ch17: 59891023-59891026) on one allele, resulting in a frameshift and premature stop after 117 amino acids, and an insertion at nt 59891024 on the other, resulting in a frameshift and premature stop after 39 amino acids. *TUBD1 -/-* line 3 is a compound heterozygote, containing a 17 nt deletion (ch17:59891015-59891031) on one allele, resulting in a frameshift and premature stop after 46 amino acids, and an insertion at nt 59891024 on the other, resulting in a frameshift and premature stop after 39 amino acids. *TUBE1 -/-* line 2 is a compound heterozygote, containing a 1049 nt deletion (ch6:112086549- 112087598) on one allele, removing exon 1 and the ATG, and a 329 nt deletion (ch6: 112087153-112087482) and 4 nt insertion (CCGA) on the other allele, removing the first exon and the ATG. The next ATG is not in-frame for any mutants. **B)** Quantification of centriole number distribution in asynchronous cells for independently-derived *TUBD1 -/-* and *TUBE1 -/-* clonal cell lines, as measured by centrin and CP110 colocalization. Bars represent the mean of three independent experiments with ≥100 cells each, error bars represent the SEM. **C)** Quantification of centriole number distribution in asynchronous cells for rescue lines, as measured by centrin and CP110 colocalization. Cell line #1 for each mutant were infected with untagged delta-tubulin or epsilon-tubulin, respectively. The rescue construct also contained monomeric Kusabira Orange kappa (mKOk) under a separate promoter. mKOk-positive cells were counted for each line. Bars represent the mean of three independent experiments with >_100 cells each, error bars represent the SEM. D) Centrin and gamma-tubulin colocalization in the indicated cell lines. Control cells are RPE-1 *TP53 -/-*. Scale bars: 5 μm.

**Figure 4 – figure supplement 1:**
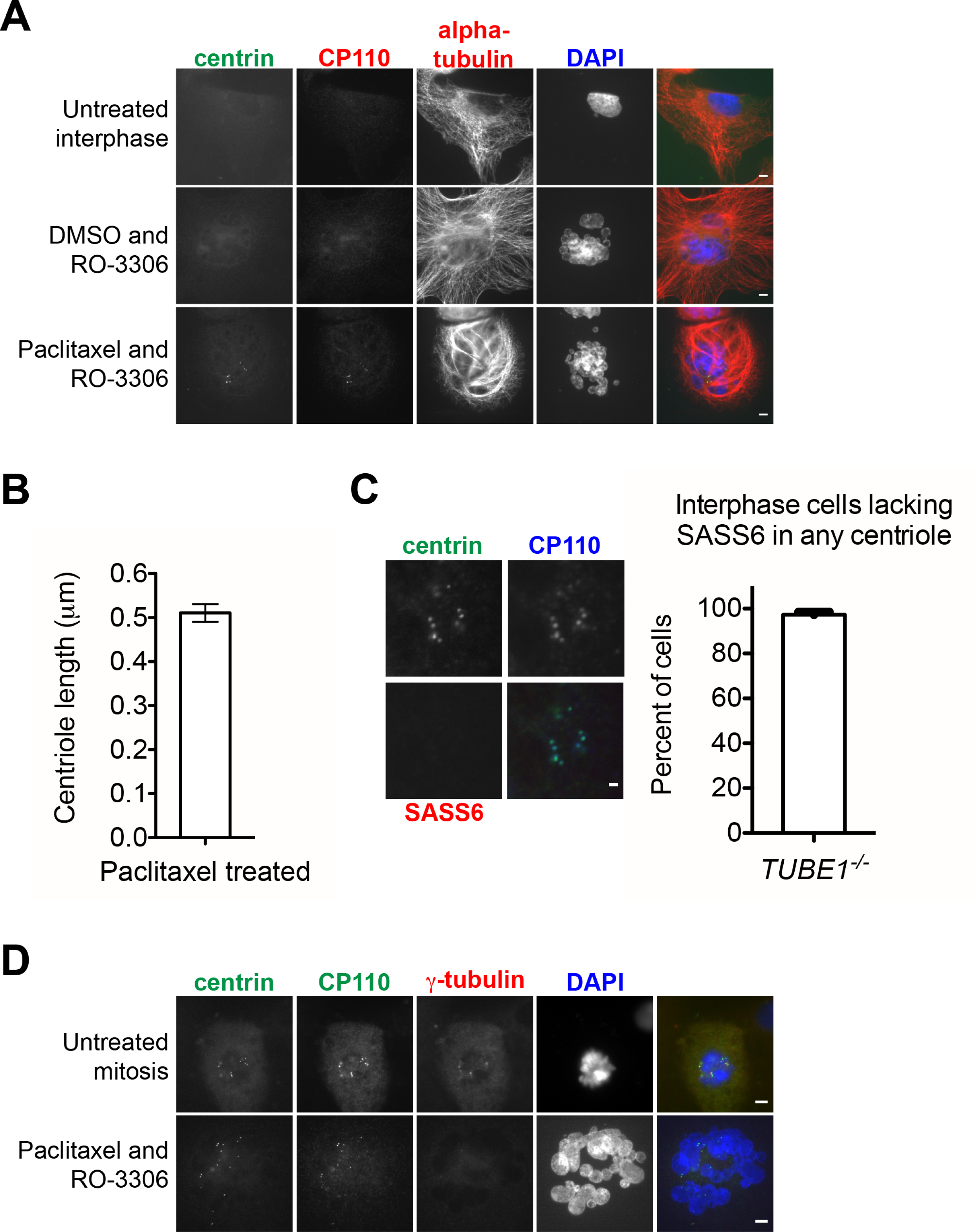
Aberrant centrioles are stabilized upon treatment with paclitaxel, despite losing SASS6 and pericentriolar material. **A)** Stabilization of *TUBE1 -/-* centrioles with paclitaxel treatment. Untreated *TUBE1 -/-* interphase cells were obtained by allowing cells to grow for 3 h after mitotic shakeoff. DMSO with RO-3306 and paclitaxel with RO-3306 treatments were performed as shown for Fig 4D on *TUBE1 -/-* cells. Bundled microtubules are present upon paclitaxel treatment, and micronuclei found in cells forced into G1 with RO-3306. In merged image, both CP110 and alpha-tubulin are represented in red. Scale bars: 5 pm. **B)** Quantification of CP110 and SASS6 separation distance in mitotic *TUBE1 -/-* cells treated with paclitaxel. 36 centrioles were measured, and measurements are not significantly different from *TUBE1 -/-* mitotic cells (compared to Fig 2C by two-tailed unpaired t-test, p=0.62). Error bars represent the SEM. **C)** SASS6 is lost in stabilized centrioles in *TUBE1 -/-* cells. Cells were treated with paclitaxel, then forced into G1 with RO-3306 as in Fig 4D. Left: Images of centrioles stained for centrin, CP110, and SASS6. Scale bar: 1 pm. Right: Quantification of percent of cells that lack any centriolar SASS6. Bars represent the mean of three independent experiments with >_100 cells each, error bars represent the SEM. **D)** Stabilized centrioles in *TUBE1 -/-* cells lack gamma-tubulin. *TUBE1 -/-* cells were either untreated and analyzed at mitosis, or treated with paclitaxel and RO-3306. Scale bars: 5 pm.

**Figure 5 – figure supplement 1:**
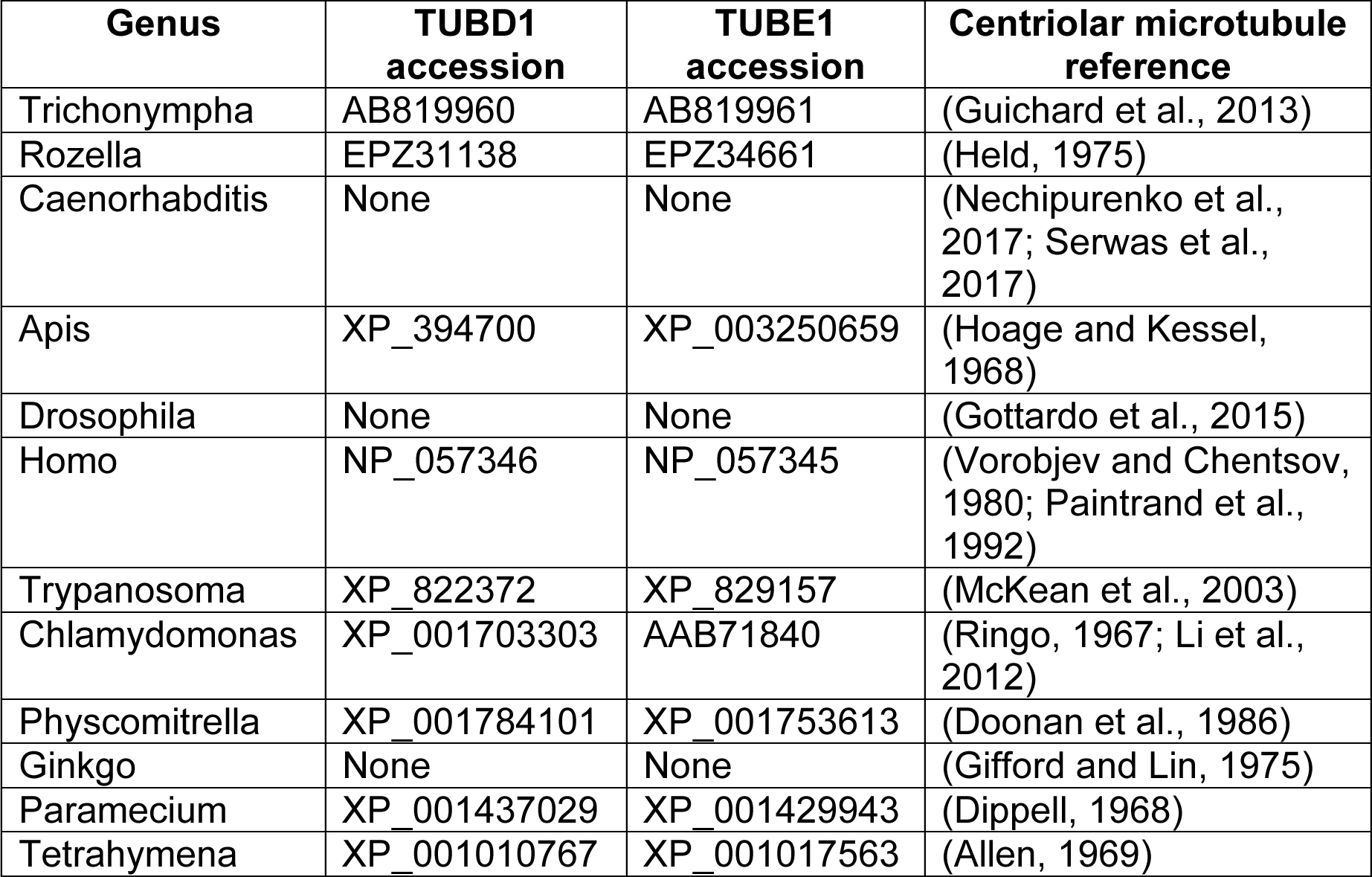
**Expanded evolutionary analysis**

## References

Allen, R.D. (1969). The morphogenesis of basal bodies and accessory structures of the cortex of the ciliated protozoan Tetrahymena pyriformis. J Cell Biol 40, 716–733.

Azimzadeh, J., Hergert, P., Delouvée, A., Euteneuer, U., Formstecher, E., Khodjakov, A., and Bornens, M. (2009). hPOC5 is a centrin-binding protein required for assembly of full-length centrioles. The Journal of Cell Biology 185, 101–114.

Balestra, F.R., Tobel, von, L., and Gónczy, P. (2015). Paternally contributed centrioles exhibit exceptional persistence in. Cell Res. 25, 642–644.

Bazzi, H., and Anderson, K.V. (2014). Acentriolar mitosis activates a p53-dependent apoptosis pathway in the mouse embryo. Proceedings of the National Academy of Sciences 111, E1491–E1500.

Cong, L., Ran, F.A., Cox, D., Lin, S., Barretto, R., Habib, N., Hsu, P.D., Wu, X., Jiang, W., Marraffini, L.A., Zhang, F. (2013). Multiplex genome engineering using CRISPR/Cas Systems. Science 339, 819–823.

Dippell, R.V. (1968). The development of basal bodies in Paramecium. Proceedings of the National Academy of Sciences 61, 461–468.

Doonan, J.H., Lloyd, C.W., and Duckett, J.G. (1986). Anti-tubulin antibodies locate the blepharoplast during spermatogenesis in the fern Platyzoma microphyllum R.Br.: a correlated immunofluorescence and electron-microscopic study. Journal of Cell Science 81, 243–265.

Dupuis-Williams, P., Fleury-Aubusson, A., de Loubresse, N.G., Geoffroy, H., Vayssié, L., Galvani, A., Espigat, A., and Rossier, J. (2002). Functional role of epsilon-tubulin in the assembly of the centriolar microtubule scaffold. J Cell Biol 158, 1183–1193.

Dutcher, S.K., and Trabuco, E.C. (1998). The UNI3 gene is required for assembly of basal bodies of Chlamydomonas and encodes delta-tubulin, a new member of the tubulin superfamily. Molecular Biology of the Cell 9, 1293–1308.

Dutcher, S.K., Morrissette, N.S., Preble, A.M., Rackley, C., and Stanga, J. (2002). Epsilon-tubulin is an essential component of the centriole. Molecular Biology of the Cell 13, 3859–3869.

Findeisen, P., Mühlhausen, S., Dempewolf, S., Hertzog, J., Zietlow, A., Carlomagno, T., and Kollmar, M. (2014). Six subgroups and extensive recent duplications characterize the evolution of the eukaryotic tubulin protein family. Genome Biol Evol 6, 2274–2288.

Fu, J., Lipinszki, Z., Rangone, H., Min, M., Mykura, C., Chao-Chu, J., Schneider, S., Dzhindzhev, N.S., Gottardo, M., Riparbelli, M.G., et al. (2015). Conserved molecular interactions in centriole-to-centrosome conversion. Nature Cell Biology.

Gadelha, C., Wickstead, B., McKean, P.G., and Gull, K. (2006). Basal body and flagellum mutants reveal a rotational constraint of the central pair microtubules in the axonemes of trypanosomes. Journal of Cell Science 119, 2405–2413.

Garreau de Loubresse, N., Ruiz, F., Beisson, J., and Klotz, C. (2001). Role of delta-tubulin and the C-tubule in assembly of Paramecium basal bodies. BMC Cell Biol. 2, 4.

Gifford, E.M., Jr, and Lin, J. (1975). Light microscope and ultrastructural studies of the male gametophyte in Ginkgo biloba: the spermatogenous cell. American Journal of Botany.

Goodenough, U.W., and StClair, H.S. (1975). BALD-2: a mutation affecting the formation of doublet and triplet sets of microtubules in Chlamydomonas reinhardtii. J Cell Biol 66, 480–491.

Gottardo, M., Callaini, G., and Riparbelli, M.G. (2015). The Drosophila centriole - conversion of doublets into triplets within the stem cell niche. Journal of Cell Science 128, 2437–2442.

Graser, S., Stierhof, Y.-D., Lavoie, S.B., Gassner, O.S., Lamla, S., Le Clech, M., and Nigg, E.A. (2007). Cep164, a novel centriole appendage protein required for primary cilium formation. J Cell Biol 179, 321–330.

Guarguaglini, G., Duncan, P.I., Stierhof, Y.D., Holmström, T., Duensing, S., and Nigg, E.A. (2005). The forkhead-associated domain protein Cep170 interacts with Polo-like kinase 1 and serves as a marker for mature centrioles. Molecular Biology of the Cell 16, 1095–1107.

Guichard, P., Hamel, V., Le Guennec, M., Banterle, N., Iacovache, I., Nemčíková, V., Flückiger, I., Goldie, K.N., Stahlberg, H., Lévy, D., et al. (2017). Cell-free reconstitution reveals centriole cartwheel assembly mechanisms. Nat Commun 8, 14813.

Guichard, P., Hachet, V., Majubu, N., Neves, A., Demurtas, D., Olieric, N., Flückiger, I., Yamada, A., Kihara, K., Nishida, Y., et al. (2013). Native architecture of the centriole proximal region reveals features underlying its 9-fold radial symmetry. Curr. Biol. 23, 1620–1628.

Held, A.A. (1975). The zoospore of Rozella allomycis: ultrastructure. Canadian Journal of Botany.

Hilbert, M., Noga, A., Frey, D., Hamel, V., Guichard, P., Kraatz, S.H.W., Pfreundschuh, M., Hosner, S., Flückiger, I., Jaussi, R., et al. (2016). SAS-6 engineering reveals interdependence between cartwheel and microtubules in determining centriole architecture. Nature Cell Biology 18, 393–403.

Hoage, T.R., and Kessel, R.G. (1968). An electron microscope study of the process of differentiation during spermatogenesis in the drone honey bee (Apis mellifera L.) with special reference to centriole replication and elimination. J. Ultrastruct. Res. 24, 6–32.

Khodjakov, A., and Rieder, C.L. (1999). The sudden recruitment of gamma-tubulin to the centrosome at the onset of mitosis and its dynamic exchange throughout the cell cycle, do not require microtubules. J Cell Biol 146, 585–596.

Kochanski, R.S., and Borisy, G.G. (1990). Mode of centriole duplication and distribution. J Cell Biol 110, 1599–1605.

Izquierdo, D., Wang, W.-J., Uryu, K., and Tsou, M.-F.B. (2014). Stabilization of cartwheel-less centrioles for duplication requires CEP295-mediated centriole-to-centrosome conversion. Cell Rep 8, 957–965.

Kleylein-Sohn, J., Westendorf, J., Le Clech, M., Habedanck, R., Stierhof, Y.-D., and Nigg, E.A. (2007). Plk4-induced centriole biogenesis in human cells. Developmental Cell 13, 190–202.

Kong, D., and Loncarek, J. (2015). Correlative light and electron microscopy analysis of the centrosome: A step-by-step protocol. Methods Cell Biol. 129, 1–18.

Lambrus, B.G., Uetake, Y., Clutario, K.M., Daggubati, V., Snyder, M., Sluder, G., and Holland, A.J. (2015). p53 protects against genome instability following centriole duplication failure. The Journal of Cell Biology 210, 63–77.

La Terra, S., English, C.N., Hergert, P., McEwen, B.F., Sluder, G., and Khodjakov, A. (2005). The de novo centriole assembly pathway in HeLa cells: cell cycle progression and centriole assembly/maturation. The Journal of Cell Biology 168, 7130722.

Lee, Y.L., Sante, J., Comerci, C., Cyge, B., and Stearns, T. (2014). Cby1 promotes Ahi1 recruitment to a ring-shaped domain at the centriole-cilium interface and facilitates proper cilium formation and function. 1–46.

Li, S., Fernandez, J.-J., Marshall, W.F., and Agard, D.A. (2012). Three-dimensional structure of basal body triplet revealed by electron cryo-tomography. Embo J. 31, 552–562.

Loncarek, J., Hergert, P., Magidson, V., and Khodjakov, A. (2008). Control of daughter centriole formation by the pericentriolar material. Nature Cell Biology 10, 322–328.

McKean, P.G., Banes, A., Vaughan, S., Gull, K. (2003). Gamma-tubulin functions in the nucleation of a discrete subset of microtubules in the eukaryotic flagellum. Current Biology 13, 598–602.

Meehl, J.B., Bayless, B.A., Giddings, T.H., Pearson, C.G., and Winey, M. (2016). Tetrahymena Poc1 ensures proper intertriplet microtubule linkages to maintain basal body integrity. Molecular Biology of the Cell 27, 2394–2403.

Nechipurenko, I.V., Berciu, C., Sengupta, P., and Nicastro, D. (2017). Centriolar remodeling underlies basal body maturation during ciliogenesis in Caenorhabditis elegans. eLife.

Nigg, E.A., Stearns, T. (2011). The centrosome cycle: Centriole biogenesis, duplication, and inherent asymmetries. Nature Cell Biology 13, 1154–1160.

Paintrand, M., Moudjou, M., Delacroix, H., and Bornens, M. (1992). Centrosome organization and centriole architecture: their sensitivity to divalent cations. J. Struct. Biol. 108, 107–128.

Ringo, D.L. (1967). Flagellar motion and fine structure of the flagellar apparatus in Chlamydomonas. J. Cell Biol. 33, 543–571.

Ross, I., Clarissa, C., Giddings, T.H., and Winey, M. (2013). ε-tubulin is essential in Tetrahymena thermophila for the assembly and stability of basal bodies. Journal of Cell Science 126, 3441–3451.

Rothbauer, U., Zolghadr, K., Muyldermans, S., Schepers, A., Cardoso, M.C., and Leonhardt, H. (2008). A versatile nanotrap for biochemical and functional studies with fluorescent fusion proteins. Molecular & Cellular Proteomics 7, 282–289.

Sanjana, N.E., Shalem, O., and Zhang, F. (2014). Improved vectors and genome-wide libraries for CRISPR screening. Nature Methods 11, 783–784.

Serwas, D., Su, T.Y., Roessler, M., Wang, S., and Dammermann, A. (2017). Centrioles initiate cilia assembly but are dispensable for maturation and maintenance in C. elegans. The Journal of Cell Biology.

Shalem, O., Sanjana, N.E., Hartenian, E., Shi, X., Scott, D.A., Mikkelsen, T.S., Heckl, D., Ebert, B.L., Root, D.E., Doench, J.G., et al. (2014). Genome-scale CRISPR-Cas9 knockout screening in human cells. Science 343, 84–87.

Sonnen, K.F., Schermelleh, L., Leonhardt, H., and Nigg, E.A. (2012). 3D-structured illumination microscopy provides novel insight into architecture of human centrosomes. Biol Open 1, 965–976.

Tsou, M.-F.B., and Stearns, T. (2006). Mechanism limiting centrosome duplication to once per cell cycle. Nature 442, 947–951.

Tsou, M.-F.B., Wang, W.-J., George, K.A., Uryu, K., Stearns, T., and Jallepalli, P.V. (2009). Polo kinase and separase regulate the mitotic licensing of centriole duplication in human cells. Developmental Cell 17, 344–354.

Turk, E., Wills, A.A., Kwon, T., Sedzinski, J., Wallingford, J.B., and Stearns, T. (2015). Zeta-Tubulin Is a Member of a Conserved Tubulin Module and Is a Component of the Centriolar Basal Foot in Multiciliated Cells. Curr. Biol. 25, 2177–2183.

Vorobjev, I.A., and Chentsov, Y.S. (1980). The ultrastructure of centriole in mammalian tissue culture cells. Cell Biol. Int. Rep. 4, 1037–1044.

Vorobjev, I.A., and Chentsov, Y.S. (1982). Centrioles in the cell cycle. I. Epithelial cells. The Journal of Cell Biology 98, 938–949.

Wang, W.-J., Acehan, D., Kao, C.-H., Jane, W.-N., Uryu, K., and Tsou, M.-F.B. (2015). De novo centriole formation in human cells is error-prone and does not require SAS-6 self-assembly. eLife 4.

Wong, Y.L., Anzola, J.V., Davis, R.L., Yoon, M., Motamedi, A., Kroll, A., Seo, C.P., Hsia, J.E., Kim, S.K., Mitchell, J.W., et al. (2015). Cell biology. Reversible centriole depletion with an inhibitor of Polo-like kinase 4. Science 348, 1155–1160.

